# Systematic evaluation of NIPT aneuploidy detection software tools with clinically validated NIPT samples

**DOI:** 10.1101/2021.05.27.445941

**Authors:** Priit Paluoja, Hindrek Teder, Amin Ardeshirdavani, Baran Bayindir, Joris Vermeesch, Andres Salumets, Kaarel Krjutškov, Priit Palta

**Affiliations:** Doctoral Programme in Population Health, Faculty of Medicine, University of Helsinki, Helsinki, Finland; Institute of Clinical Medicine, University of Tartu, Tartu, Estonia; Competence Centre for Health Technologies, Tartu, Estonia; Institute of Biomedicine and Translational Medicine, Department of Biomedicine, University of Tartu, Tartu, Estonia; Medpace Reference Laboratories, Molecular Unit, Leuven, Belgium; Department of Human Genetics, KU Leuven, Leuven, Belgium; Department of Obstetrics and Gynaecology, Institute of Clinical Medicine, University of Tartu, Tartu, Estonia; Estonian Genome Center, Institute of Genomics, University of Tartu, Tartu, Estonia; Institute for Molecular Medicine Finland, FIMM, HiLIFE, University of Helsinki, Helsinki, Finland

## Abstract

**Motivation:** Non-invasive prenatal testing (NIPT) is a powerful screening method for fetal aneuploidy detection, relying on laboratory and computational analysis of cell-free DNA. Although several published computational NIPT analysis tools are available, no comprehensive and direct accuracy comparison of these tools is published. Here, we evaluate and determine the precision of five commonly used computational NIPT aneuploidy analysis tools, considering diverse sequencing depth (coverage) and fetal DNA fraction (FF) on clinically validated NIPT samples.

**Methods:** We evaluated computational NIPT aneuploidy analysis tools WisecondorX, NIPTeR, NIPTmer, RAPIDR, and GIPseq, on the same set of clinically validated samples, subsampled to different sequencing coverages between 1.25–20M reads per sample (RPS). These clinically validated samples consisted of 423 samples, including 19 samples with fetal chromosome 21 trisomy (T21, Down syndrome), eight trisomy 18 (T18, Edwards syndrome) and three trisomy 13 (T13, Patau syndrome) samples. For each software and sequencing coverage, we determined the number of false-negative and false-positive trisomy/euploidy calls. For a uniform trisomy detection interpretation, we defined a framework based on the percent-point function for determining the cut-off threshold for calling aneuploidy based on the sample Z-score and the reference group Z-score distribution. We also determined the effect of the naturally occurring arbitrary read placement driven uncertainty on T21 detection at very low sequencing coverage and the effect of cell-free fetal DNA fraction (FF) on the accuracy of these computational tools in the case of various sequencing coverages.

**Results:** This is the first head-to-head comparison of NIPT aneuploidy detection tools for the low-coverage whole-genome sequencing approach. We determined that, with the currently available software tools, the minimum sequencing coverage with no false-negative trisomic cases was 5M RPS. Secondly, for these compared tools, the number of false-negative trisomic cases could be reduced if the trisomy call cut-off threshold considers the Z-score distribution of euploid reference samples. Thirdly, we observed that in the case of low FF, both aneuploidy Z-score and FF inference was considerably less accurate, especially in NIPT assays with 5M RPS or lower coverage.

**Conclusions:** We determined that all compared computational NIPT tools were affected by lower sequencing depth, resulting in systematically increasing the proportions of false-negative trisomy results as the sequencing depth decreased. Trisomy detection for lower coverage NIPT samples (e.g. 2.5M RPS) is technically possible but can increase the proportion of false-positive and false-negative trisomic cases, especially in the case of low FF.

## Introduction

Non-invasive prenatal testing (NIPT) is widely used as a highly accurate screening test of fetal chromosomal aneuploidies (1). As it relies on whole-genome sequencing (WGS) or targeted sequencing of cell-free DNA (cfDNA) extracted from the peripheral blood of a pregnant woman, it reduces the number of invasive amniocentesis procedures (2, 3). NIPT analysis is completed with specific computational analyses that make use of generated WGS data to infer the excess or deficiency of sequencing reads belonging to a specific chromosome, indicating a possible aneuploidy of the corresponding chromosome in cell-free fetal DNA and fetus (3, 4).

To date, several computational NIPT analysis tools for WGS-based NIPT have been published. These include GIPseq (3), NIPTmer (5), NIPTeR (6), RAPIDR (7), DASAF R (8), Wisecondor (9), and WisecondorX (9, 10). However, while these computational tools are widely used, no head-to-head evaluation of these NIPT tools on the same clinically validated samples is available.

Computational NIPT studies illustrate that the most relevant aspects of NIPT analyses is sequencing depth (i.e. sequencing read coverage) (10, 11). The higher the sequencing depth, the more comprehensively the entire genome is interrogated, and the more evidence there is to determine any consecutive chromosome-spanning gains or losses and, ultimately, possible aneuploidies in studied samples (11). Currently, 10M reads per sample are considered reliable for detecting Down, Edwards, and Patau syndromes in the case of clinical applications (11).

A second relevant aspect of the computational NIPT aneuploidy detection is the analytical interpretation of the computational tool output. Commonly used NIPT tools output a per chromosome metric describing the difference (or similarity) of the sample of interest compared to reference group samples, representing the NIPT data of known/validated euploid samples (3, 5, 6, 9, 10). While some tools do provide explicit guidelines for interpreting the output (6), in general, the NIPT software output and analytical interpretation are not well standardised and tend to be highly dependent of the software, laboratory protocols, sample pre-processing and also reference group utilised in the process. Therefore, an universally usable framework for the interpretation of these metrics is needed.

Finally, it is relevant to consider that in the NIPT data analysis, sequencing reads originate both from the studied fetus/placenta and the mother (12). The maternal chromosomal status can be considered as a baseline and the fetal chromosomal status as the signal of interest (13). The proportion of fetal DNA fraction (FF), generating the signal of interest, is a critical sample-level quality control determinant, allowing to detect samples with too low FF, for which the computational aneuploidy calling nor euploidy confirmation cannot be performed confidently. As low-coverage WGS-based NIPT assays and the following computational analyses mostly do not distinguish between fetal and maternally originating sequencing reads (but analyse them as a whole), a sufficient proportion of fetal DNA fraction is crucial (3, 5, 6, 9, 10, 13).

In this study, we have assessed these critical aspects of NIPT and their effect on the accuracy of five widely used NIPT software tools. We systematically evaluate the published computational NIPT tools’ performance and accuracy on a set of clinically validated samples, considering various sequencing depths and the proportion of cell-free fetal DNA. We define and validate a straightforward and universal Z-score quantile cut-off based framework that can be unambiguously used to describe and compare aneuploidy calling software tools.

## Materials and Methods

### Studied samples

Two sets of samples were used. **For the reference sample set**, a total of 669 known euploid samples were used. Of those, 326 of these samples were of female fetus pregnancies and 343 of male fetus pregnancies. All 669 samples had been reported previously to be euploid by the NIPTIFY screening test. **The validation sample set** was based on a previously published validation study by Žilina *et al*. (14), consisting of 423 samples, of which 258 were high-risk pregnancies that had undergone diagnostic invasive prenatal analysis (14). These included 19 samples with confirmed fetal T21, eight T18 and three T13 samples.

All samples were sequenced with Illumina NextSeq 500 platform, producing 85 bp single-end reads with an average per-sample coverage of 0.32× at the University of Tartu, Institute of Genomics Core Facility, according to the manufacturer’s standard protocols, as described previously (14). This study was performed with the informed consent from the participants and with approval of the Research Ethics Committee of the University of Tartu (#315/T-13).

### Sample pre-processing

For the alignment-based computational NIPT methods, each sequenced sample was aligned against human reference assembly GRCh37 (RAPIDR) or GRCh38 (NIPTeR, WisecondorX) depending on the software prerequisite. Next, the aligned sample was sorted, and the reads originating from a single fragment of DNA were marked as duplicates. For the k-mer based computational workflow approach, no special pre-processing was applied.

Both the validation set and the reference set were emulated to lower sequencing coverage. For this, each sample was subsampled into six different groups of subsamples. The average million reads per sample targets were: 20M, 15M, 10M, 5M, 2.5M, and 1.25M. For 5-20M, the lower sequencing coverage was emulated by leaving the appropriate number of NextSeq 500 output lanes out. For lower and equal to 2.5M, one lane (5M) was taken and subsampled with samtools view and then converted to FASTQ with samtools bam2fq (15), and the exact resulting coverage was then calculated for each sample. Clinically validated T21, T18, and T13 sample group information with the reference population and low-risk validation sample group are presented in **Supplementary Table 1**.

### Study approach

Computational NIPT methods applied in this study use a euploid reference set. For each analysed sequencing coverage, a corresponding reference set was created. For example, for the 5M RPS coverage, both the validation and reference samples were subsampled to 5M RPS analysed as a 5M RPS group. For each sample, coverage, fetal DNA fraction and Z-score estimates for chromosomes 13, 18, and 21 were calculated.

#### Sequencing depth effect of aneuploidy detection

To analyse the effect of sequencing depth (read coverage) on aneuploidy detection, we systematically subsampled raw sequencing read data from the average of 20M reads to 1.25M reads per sample. Next, we applied a computational NIPT tool (with the reference corresponding to the RPS) to infer sample Z-scores. Then, we calculated the accuracy corresponding to different sequencing coverage for each software by counting the number of correctly or incorrectly detected known trisomies and euploid samples by comparing the sample Z-score with the empirically calculated cut-off *Z*_*e*_ (except for GIPseq, which was evaluated by the interpretations received from the GIPseq authors). We also found that NIPTeR, NIPTeR NVC, and RAPIDR do not provide results if the sequencing coverage is lower than 1.25 M reads per sample.

#### Low sequencing coverage driven uncertainty in Z-score inference for trisomic and euploid samples

We used ten low-risk samples to determine how low sequencing depth and consequent arbitrary sequencing read distribution and binning affect the uncertainty in Z-score estimation. These samples were selected to have a high fetal fraction (FF estimates between 10.23–18.57%) and high sequencing read counts (read count between 22M–30M) and with chromosome 21 Z-scores close to zero (original read coverage NIPTeR Z-scores between -0.0996 and 0.0942). The BAM files of those samples were concatenated and sorted, leading to a single pooled low-risk sample with 247M reads. The concatenated sample was then randomly subsampled 2,000 times to groups of 2.5, 5, 10 and 20M RPS, followed by the NIPTeR Z-score calculations. The expected Z-score of each generated sample is approximately zero. Next, the deviations were found by subtracting the average group Z-score from the calculated Z-scores. These normally occurring deviations from the expected Z-score of 0 were added to the original validation sample Z-scores, leading to 2,358,000 simulated low-risk Z-scores and FP and TN count (based on *Z*_*t*_). Similarly to low-risk samples, the same methodology was also applied to T21 samples. The concatenation was done with eight T21 samples (read count 20M–33M, FF 10.43%–14.21%, chromosome 21 NIPTeR Z-score 13.77–23.64) that led to 203M RPS. For this simulation, 6, 7, 8 and 9M RPS reference group was created and similarly to 1.25 and 2.5M RPS, 20M RPS samples were downsampled.

#### Sequencing depth effect on fetal DNA fraction estimation

To analyse the effect of sequencing depth on FF estimation, we first estimated FF for the 20M RPS group and divided samples into FF groups of 0–5% and 5–15% based on the estimated FF. Next, we systematically subsampled raw sequencing read data from the average of 20M reads to 1.25M reads per sample. Finally, we compared the FF estimates of different sequencing coverages (also considering the FF group) with 20M by calculating Pearson correlation.

### Aneuploidy detection with empirically defined Z-score thresholds

All evaluated computational NIPT tools provided Z-score output for assessing the risk for aneuploidy. Each of the algorithms also has specific differences. For example, RAPIDR outputs trisomy calls (7), WisecondorX creates log_2_ ratio chromosome figures between the ratio of the observed number and expected number of reads (10), NIPTmer publication defines cut-off at Z-score of 3.5 (5), GIPseq provides algorithmic decision tree-like interpretation of the results (3), and NIPTeR avoids defining clear trisomy call cut-off (6). However, to compare different algorithms’ performance on similar grounds, we defined a generally usable framework for Z-score calculations and comparisons, relying on a straightforward percent point (quantile) function. Specific quantile point allows specifying the cut-off value of the computational NIPT Z-score such that the probability of the euploid sample Z-score being less than or equal to the cut-off equals the chosen probability. This calculation can be done with the presumption that the mean and SD of the euploid sample group Z-score distribution is 0 and 1 (standard normal distribution) or with empirically observed mean and SD of the euploid sample reference group Z-score distribution. The first threshold is referenced as a (universal) theoretical cut-off *Z*_*t*_ in the calculations and the latter as the empirical cut-off *Z*_*e*_. A probability of 0.9999 was used for all the performed analyses: P_99.99_ (P[Z_sample_ ≤ z_cut off threshold_]) = 0.9999. Theoretically, given an informative NIPT assay data with sufficient coverage and a fetal fraction, it is expected to get one false-positive trisomy call per every 10,000 analysed normal euploid samples.

### Compared software tools

For a comparison between different tools, the Z-score was used for scoring as all the computational NIPT tools supported it in the analysis. Instead of the older Wisecondor software, we decided to use the newer WisecondorX. Also, DASAF R (8) is made available by the authors, but the corresponding web links in the publication to the software are inoperable, and the software was not included. Execution and the analysis of the output of the tools were done by scripting in WDL, Python, R, and Bash in the computer cluster of the High Performance Computing Center of the University of Tartu (16). If the tool failed to operate on low coverage, then analysis of that coverage for the failed tool was left out. The most relevant data-analysis procedures, parameters, and analysis aspects for each software are shortly described and discussed in the paragraphs below.

#### NIPTeR

For NIPTeR, v1.0.2 was used. In the reference group’s creation, each sample was binned with *bin_bam_sample* (parameter *separate_strands* set to *TRUE*), then GC corrected (method *gc_correct*, parameters *method* set to *‘bin’, ref_genome* set to *‘hg38’*, and *include_XY* set to *FALSE*). After that, all the GC-corrected samples were marked as the control group. No control group matching was done as all the tested tools had to have the same control group. For each validation sample, the sample was binned and GC corrected. For scoring, each sample was chi corrected (*chi_correct(sample, control_group)*), and after that, the Z-score was calculated with the function *calculate_z_score*. Additionally, since NIPTeR supports normalized chromosome value (NIPTeR NCV), NCV was calculated with *prepare_ncv(max_elements = 9)* and *calculate_ncv_score*.

#### WisecondorX

Wisecondor (Paco_0.1) was tested on a 5M RPS group using the quick start guide published on their official GitHub page. Due to the Wisecondor output not containing a single Z-score for the chromosome but multiple Z-scores for the chromosome regions, Wisecondor was left out from the further analysis due to the incomparability with the other tested software metrics and the existence of WisecondorX, which provides a Z-score for the entire chromosome. For WisecondorX, v1.1.5 was used. Pre-processed samples were converted to .npz format with *WisecondorX convert* command. Next, the reference was created with *WisecondorX newref --nipt --refsize 669 --binsize 100000* directive. For Z-scores, the *chr_statistics* file from the output was used after applying the *predict* command.

#### NIPTmer

NIPTmer binaries were obtained from the University of Tartu Department of Bioinformatics webpage. Pre-built lists, which were packaged with the software, were used. Although the cleaned lists were made with the GRCh37 reference genome, since it is not an alignment-based method and GRCh37 lists are known to work (14), the GRCh37 version was kept. Binaries were updated to the latest version due to software issues with the packaged binary.

#### GIPseq

The GIPseq (3) was run at the KU Leuven. The samples were uploaded to the KU Leuven Google Cloud bucket and analysed by GIPseq. The raw output of the analysis was shared with the authors. The GIPseq NIPT pipeline provides a sophisticated output, including quality scores and analysability for each sample. They also define specific decision rules for calling each sample state and have more states than euploid or aneuploid, including monosomy, segmental or undetermined. Since most of the other computational NIPT tools used in the analysis (I) provide only Z-score output, (II) do not have a pre-defined cut-off threshold, (III) do not provide more scores than Z-scores, the GIPseq pipeline is not directly comparable with other computational NIPT tools. For assessing the number of false-negatives and false-positives and the effect of fetal fraction, GIPseq authors interpretations (euploid or trisomy) for each sample were used. For assessing the effect of the average sample read count for Z-scores, the theoretical Z-score cut-off was used for comparability between different computational tools.

#### RAPIDR

For RAPIDR, v0.1.1 was used. Since RAPIDR requires according to the manual GenomicRanges of version 1.14.4, which is challenging to acquire due to its old release date and RAPIDR will not work with later versions, a workaround was found, allowing to use RAPIDR 0.1.1 with most recent R packages. For this, R package GenomicAlignments (bioconductor-genomicalignments 1.22.0) was installed and loaded before loading RAPIDR. GenomicAlignments addressed the issue of the missing function *summarizeOverlaps* from GenomicRanges (bioconductor-genomicranges 1.38.0). Similar to SeqFF, RAPIDR only works with the GRCh37 human reference genome. The corresponding author listed in the CRAN was enquired if and how it would be possible to run RAPIDR with the hg38 reference genome. No response was received. In terms of reference creation set, the function *makeGCContentData* was used to calculate the GC content information for the GRCh37 reference. Next, the reference samples were binned using *makeBinnedCountsFile* with parameter *k* (bin size) set to *20000* (default). The samples were binned with *makeBinnedCountsFile*, and the final R object representing the reference set was made with *createReferenceSetFromCounts* with *gcCorrect* and *filterBin* set to *TRUE*. RAPIDR does output trisomy calls in addition to Z-scores, but this calling is based on the fixed Z-score cut-off threshold of three (7). However, for uniform trisomy detection comparison over computational tools, a Z_e_ was used.

### Fetal DNA fraction calculation

For the original sequencing data and subsampled lower coverage sample sets, cell-free fetal DNA fraction (FF) was calculated with the SeqFF software (17). For SeqFF, all analysed samples were aligned against the human reference assembly GRCh37 and filtered by the alignment quality of Q30. After the filtering, the reads were sorted, and SeqFF estimates were calculated.

## Results

We compared the results of five NIPT computational software tools on the same set of clinically validated NIPT samples. By systematically subsampling sequencing reads of studied samples (to artificially lower their sequencing coverage) and by analysing the chromosomal Z-scores obtained with different NIPT software tools, we determined their ability to detect known trisomies and confirm euploid samples. We also determined the minimal sequencing depth for each NIPT software while considering fetal DNA fraction (FF) in analysed samples.

### Sequencing depth effect on aneuploidy detection

First, we calculated the number of true and false trisomy findings for each software at different analysed sequencing depths by considering the same uniform empirically defined Z-score Z_e_ threshold for each NIPT software (see Methods). Corresponding results for chromosome 21 trisomy (T21) detection accuracy are presented in **Figure 1**, and corresponding trisomy 13 (T13) and trisomy 18 (T18) detection accuracy results are presented in **Supplementary Figure 1** and **Supplementary Figure 2**.

**Figure 1.**
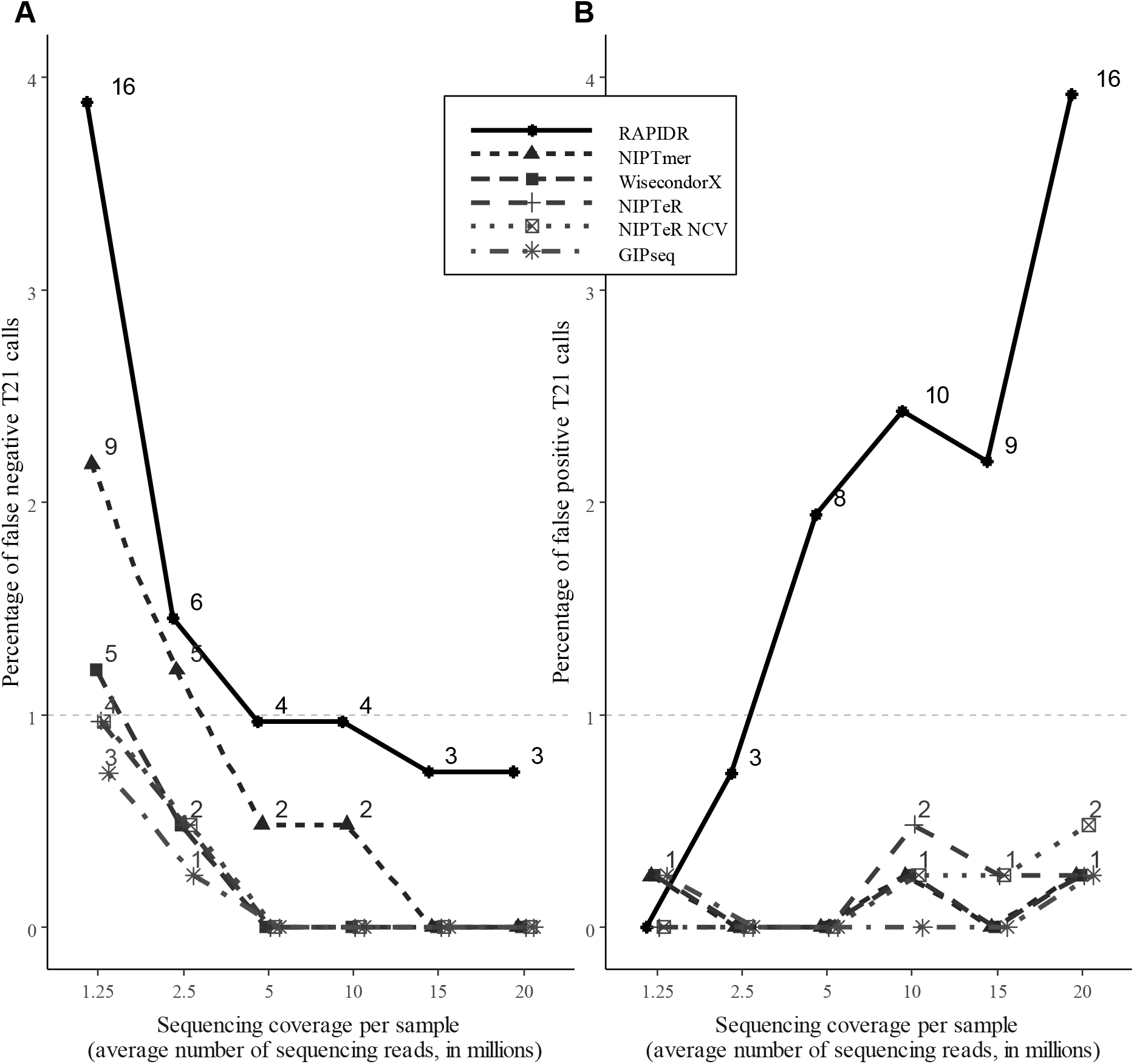
Trisomy detection accuracy of tested NIPT software tools in case of different sequencing depths. Percentage and the absolute number of (A) false-negative trisomy 21 (T21) and (B) false-positive T21 calls obtained with each NIPT software tool in case of various sequencing coverages.

Although mostly accurate with sequencing coverages higher than 5M RPS, clear trends were observed for compared algorithms in the case of lower sequencing depths. We observed substantial differences between different tools considering both false-positive and false-negative trisomy calls. Furthermore, all algorithms demonstrated a considerable increase in the number of false-negative trisomy findings below 5M RPS (**Figure 1** and **Supplementary Table 2**). These changes in trisomy detection accuracy were driven by more conservatively estimated Z-scores, systematically decreasing at lower sequencing depths (**Figure 2**).

**Figure 2.**
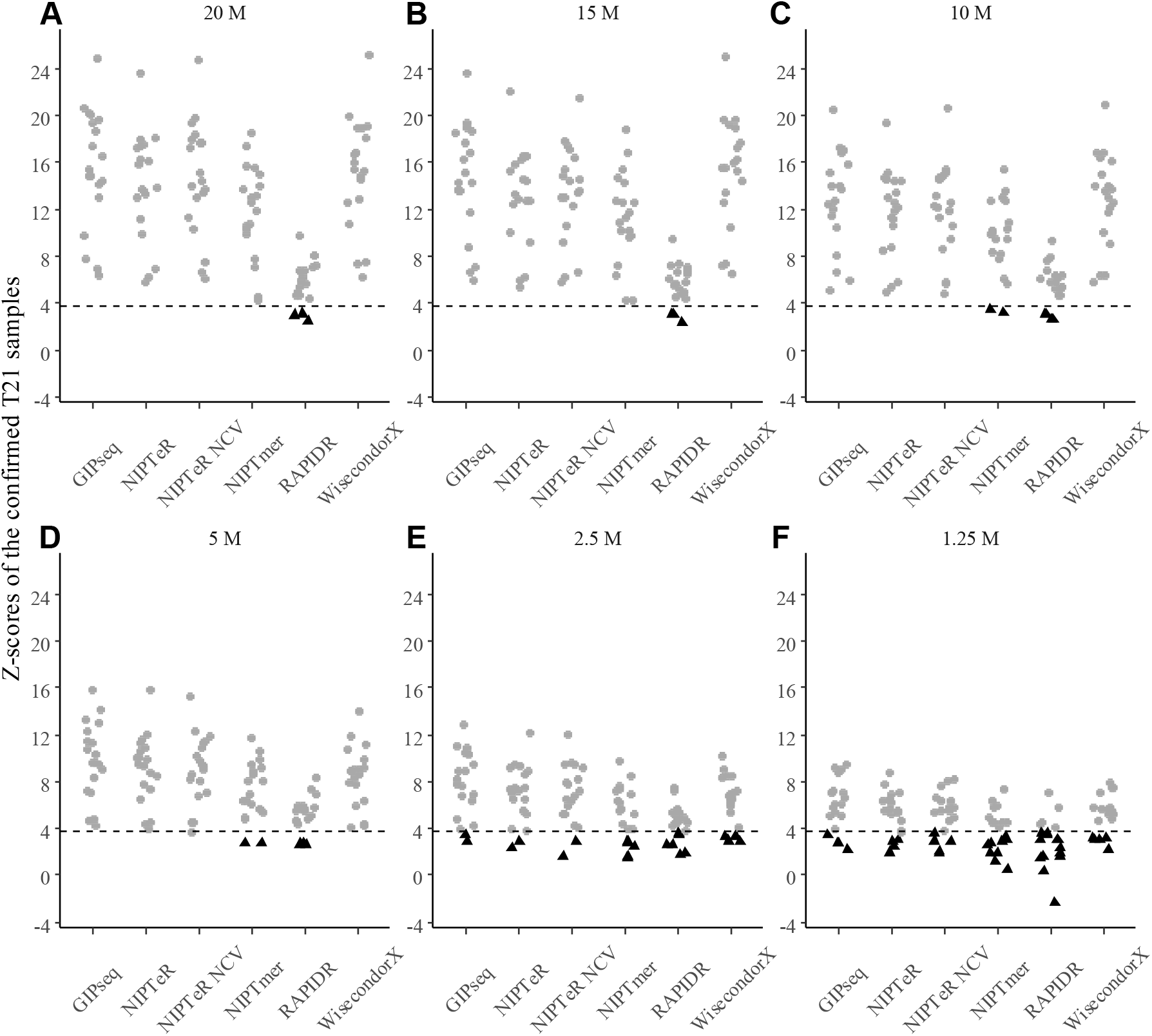
Z-scores of known clinically validated trisomy 21 samples at different sequencing depths. Z-scores of known trisomy samples in the case of sequencing depths of 20M RPS (A), 15M RPS (B), 10M RPS (C), 5M RPS (D), 2.5M RPS (E) and 1.25M RPS (F). False-negative (FN) trisomies are represented as black triangles, determined by the *Z*_*t*_ cut-off threshold (the black dashed line).

Interestingly, chromosome-specific accuracy differences were observed in the case of NIPTeR and NIPTeR-NCV. Although both detected T21 cases equally accurately, in T18 detection analyses, NIPTeR-NCV showed slightly higher accuracy, especially with regard to lower sequencing coverage.

In our comparison, GIPseq, NIPTeR and WisecondorX performed similarly well for sequencing depths of 10M RPS and higher, followed by NIPTmer and RAPIDR. The GIPseq, WisecondorX, and NIPTeR algorithms also work equally accurately at sequencing coverages lower than 10M RPS and produced very similar results in T21 and T18 detection. Furthermore, our results demonstrate that the latter tools can be relatively accurately used for trisomy detection even in the case of extremely low sequencing coverages (e.g. 2.5M and 1.25M RPS).

The above results clearly demonstrate that sufficient sequencing coverage is required for accurate NIPT aneuploidy inference. Besides considering how lower sequencing depth alters the reference panel driven changes in Z-score variability and increased uncertainty (especially at lower coverages), it is also relevant to consider how lower sequencing depth affects naturally occurring arbitrary sequencing read placement and consequent uncertainty in the studied sample’s Z-score estimation. In the corresponding analyses (see Methods), we observed that at lower sequencing coverage (7M RPS and below), the arbitrary sequencing read placement considerably influences trisomy calling accuracy. The corresponding simulations with clinically confirmed trisomy cases (**Supplementary Figure 3**) and euploid samples (**Supplementary Figure 4**) demonstrate that if the aneuploidy calling algorithm does not consider this additional source of uncertainty (especially at sequencing depths below 7M RPS), a proportion of trisomy cases can easily be undetected. Therefore, to confidently use very low sequencing coverage, appropriate correction methods are required for computational NIPT analytical tools.

While working with the Z-scores for the euploid reference group (which were used to determine the empirical trisomy calling cut-off of *Z*_*e*_), we also observed that the empirical *Z*_*e*_ and the theoretical *Z*_*t*_ cut-off thresholds are not always equal (**Supplementary Figure 5**). Importantly, these cut-offs directly affect the number of false-negative trisomy cases, e.g. for WisecondorX, the *Z*_*e*_ threshold for chromosome 21 varied from 3.14 to 4.16, depending on the used sequencing coverage. With the theoretical cut-off *Z*_*t*_, WisecondorX introduces two additional (total of four) undetected trisomies at the 2.5M RPS sequencing coverage (**Figure 1** and **Figure 2E**), as compared to the hereby suggested usage of empirically derived (reference data variance aware) threshold of *Z*_*e*_. This observation allows us to believe that NIPT software tools can benefit from *Z*_*e*_ cut-off calibration as a universal uniform cut-off *Z*_*t*_ does not consider differences between various NIPT tools and analysed data (which can also harbour laboratory-specific effects), leading to false-negative and false-positive trisomy predictions.

### Accuracy of fetal DNA fraction estimation in case of lower sequencing depths

Ideally, the FF estimate would not change for the same sample, even at the lower sequencing coverages. While considering the FF fraction estimated for each sample with the highest sequencing depth (∼20M RPS) as the ‘truth’ (see Methods), we observed that FF estimates were inconsistent at the lower sequencing depths (**Figure 3** and **Supplementary Figure 6**). In the true FF range of 0–5%, the Pearson correlation between 1.25M RPS FF estimates and the corresponding 20M RPS FF estimates was only 0.217. For the same samples, the FF correlations were relatively consistent at the higher sequencing depths, e.g. the correlation between 10M RPS FF and 20M RPS FF estimates was 0.841. For higher FF values (with true FF of 5–15%), these differences were more subtle. For 1.25M RPS FF and 20M RPS FF estimates, we observed a correlation of 0.636 and the correlation between 10M RPS FF and 20M RPS FF estimates was 0.959 (**Supplementary Figure 6**).

**Figure 3.**
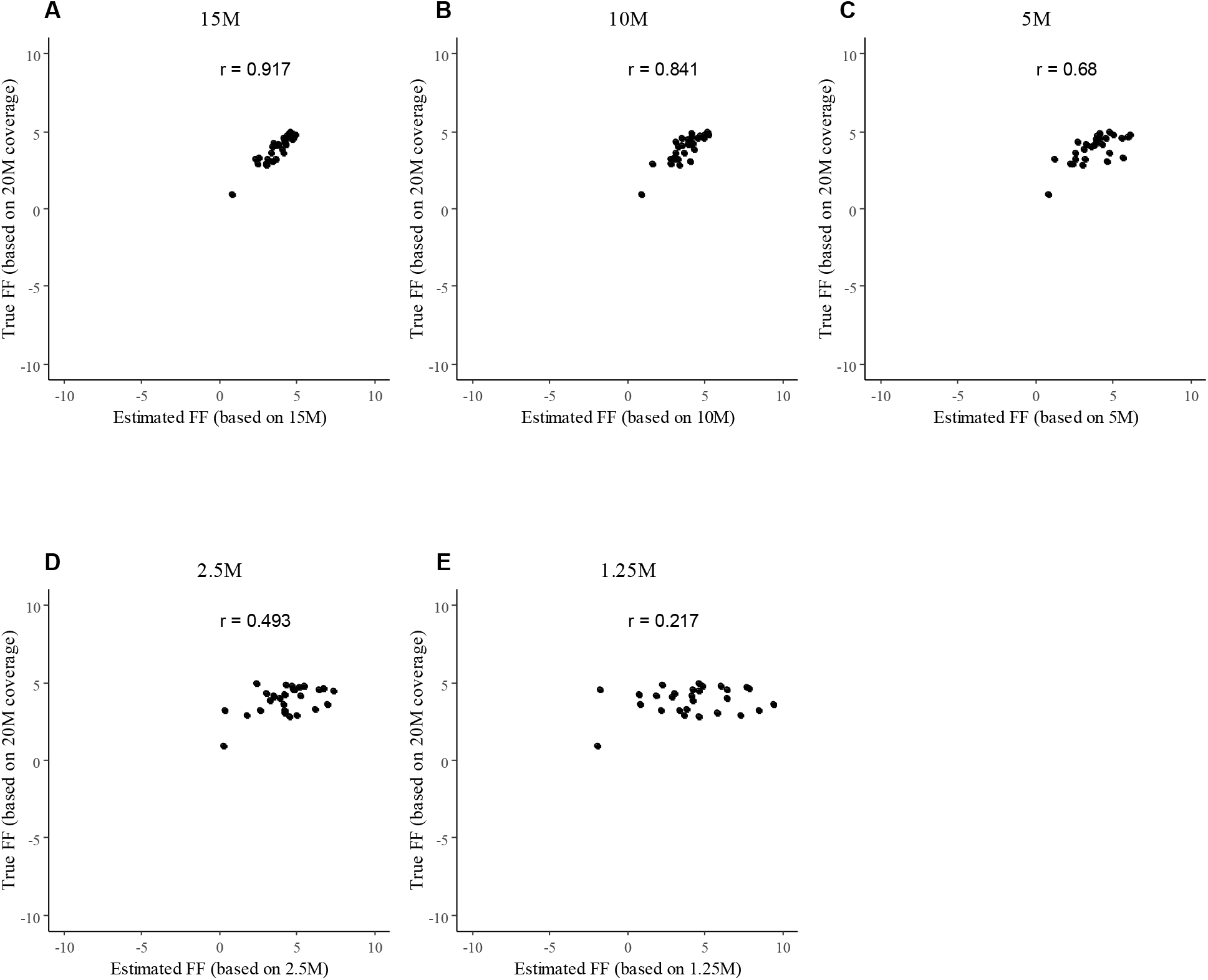
Correlation between the ‘true’ (based on 20M RPS data) and lower sequencing coverage based FF estimates. The Pearson correlation of the 20M RPS FF 0–5% estimates and the estimates on sequencing depths of 15M RPS (a), 10M RPS (b), 5M RPS (c), 2.5M RPS (d), and 1.25M RPS (e).

These patterns of correlations between the different FF estimates at the lower sequencing depths for FF of 0–5% and 5–15% suggest that FF is systematically incorrectly estimated for lower RPS samples. Such inconsistent estimates suggest a risk of flawed ‘overconfidence’ when determining aneuploidy/euploidy status at lower sequencing depths. For example, a scenario where a studied sample with a very low fetal DNA fraction (which should, in clinical applications, fail the FF-based quality control) is assayed at a low (e.g. ∼5M RPS) sequencing coverage. Since the FF of such sample can be consequently mistakenly estimated as sufficiently high, this sample would be analysed for aneuploidies and the resulting decision for chromosomal euploidy or trisomy would be erroneously considered as a confident one. At the lower sequencing depths, our further analysis confirmed this possibility (**Figure 4, Supplementary Figure 7, Supplementary Figure 8**). Notably, in the case of the lowest sequencing depth (1.25M RPS), all samples with FF below 7% (with one exception in the case of GIPseq) were undetected and incorrectly classified as normal euploid samples (**Figure 4F**). At 2.5M RPS, all tested software tools failed to detect at least one (out of 19) known T21 sample (**Figure 4E**). For 5M RPS and higher coverages, most software tools were able to detect T21 samples correctly, with only exceptions being NIPTmer and RAPIDR with two and four (out of 19) undetected trisomies in the case of 5M RPS, respectively.

**Figure 4.**
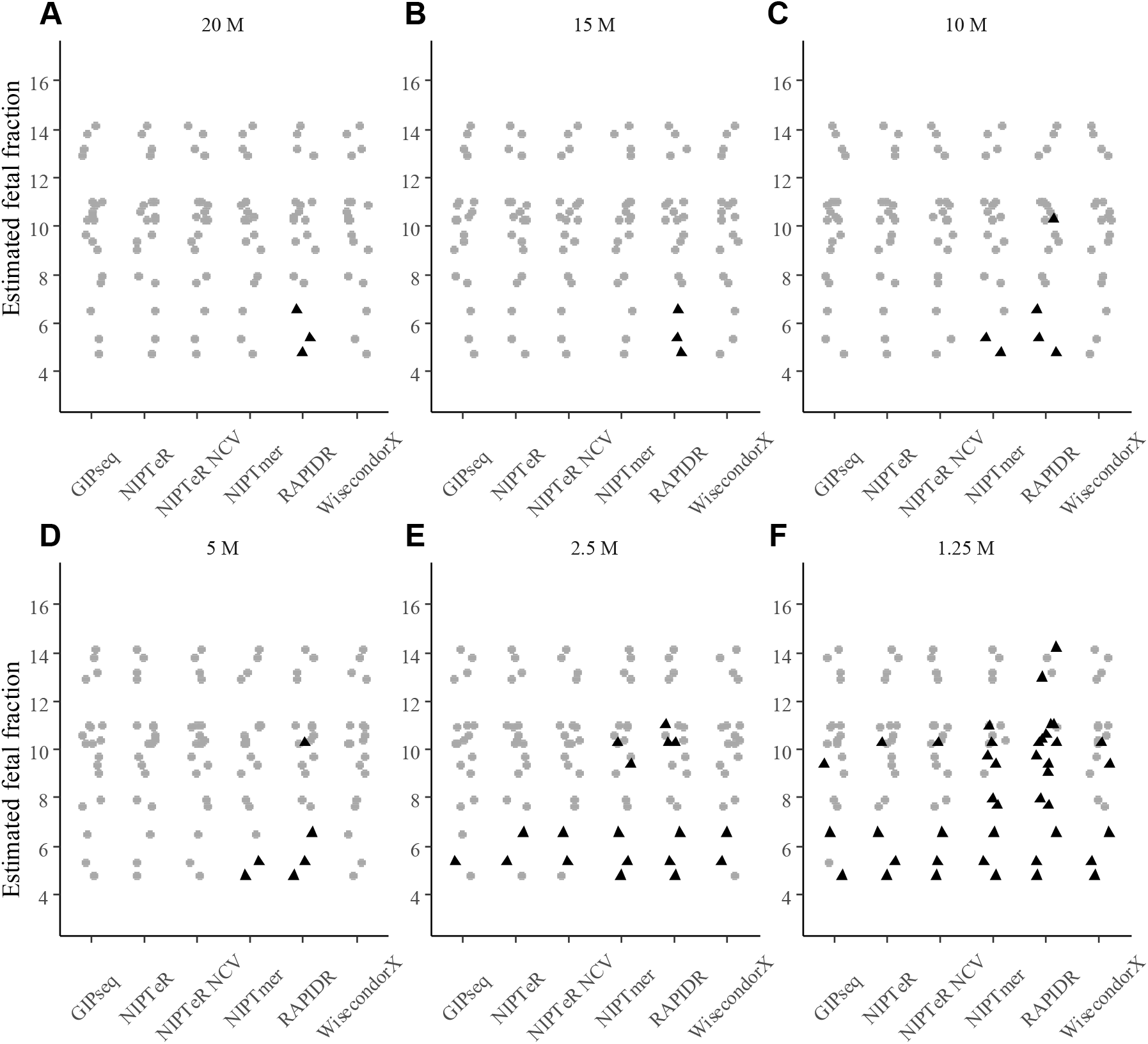
The effect of fetal DNA fraction on detecting known chromosome 21 trisomy samples at different sequencing coverages. Z_e_ cut-off was used for calling trisomy (internal classification in the case of GIPseq). Black triangles represent undetected trisomy cases. The fetal fraction calculation is based on the 20M RPS group. Sequencing depths of 20M RPS (A), 15M RPS (B), 10M RPS (C), 5M RPS (D), 2.5M RPS (E) and 1.25M RPS (F) were used.

Interestingly, GIPseq resulted in one undetected T21 case at 2.5M RPS that is still detected on lower 1.25M RPS (**Figure 4E, Figure 4F**). At the same time, there is another 1.25M RPS sample detected by GIPseq but not with any other software (**Figure 4F**). The reason for such behaviour is unknown but allows believing that it could be from the arbitrary sequencing read placement.

To conclude, our findings demonstrate that accurate trisomy detection with WGS-based NIPT is mainly affected by the computational tool selected in combination with sufficient sequencing coverage and accurately estimated fetal DNA fraction.

## Discussion

Non-invasive prenatal testing (NIPT) is an effective screening method for fetal aneuploidy testing, which is based on laboratory and computational analysis of cell-free DNA derived from the peripheral blood of pregnant women. Hence, a correctly set up NIPT assay allows reducing invasive procedures while still enabling detection of fetal aneuploidies at high confidence. Although there are several computational NIPT tools for 2^nd^ generation WGS-based NIPT, there are no published comparisons of these tools, which would allow NIPT laboratories to select an optimal analytical software matching their NIPT sequencing assays.

As expected, we found that the currently publicly available computational tools can accurately detect chromosome 13 trisomy (causing Patau syndrome), trisomy 18 (Edwards) and trisomy 21 (Down) at 5M RPS and higher sequencing coverages. We also observed that a well-chosen computational NIPT software in combination with appropriately applied Z-score thresholds can be used at lower sequencing coverages, even below 5M RPS. This is in line with the current knowledge, also NIPT potential at lower coverages (such 2.2M RPS) has been shown (18). However, at lower sequencing coverages (below 5M RPS), various computational tools infer trisomy risks with a quite different accuracy, mostly with a systematically increasing number of false-positive and false-negative trisomy cases (**Figure 1, Supplementary Figure 1, Supplementary Figure 2**). Furthermore, at very low sequencing coverages, it is also relevant to consider naturally occurring arbitrary sequencing read placement and the resulting uncertainty in the studied sample’s Z-score inference. If not accounted for, this phenomenon can lead to undetected trisomies (**Supplementary Figure 3**).

We also determined that applied algorithms are differently affected by very low fetal DNA fraction (**Figure 4, Supplementary Figure 7, Supplementary Figure 8**). In general, low sequencing coverage coupled with low FF further decreases trisomy detection accuracy (**Figure 4, Supplementary Figure 8**).

That said, there are NIPT algorithms that work accurately with FF lower than 4% (19). However, Koc *et al*. also observed that NIPT test failures (as no result) are often related to the low FF estimation in the case of lower sequencing coverage (18). Also, depending on the software used for FF calculation, FF estimator (17) accuracy can significantly decrease at low sequencing coverages in the case of truly low FF (**Figure 3, Supplementary Figure 6**). Similar results were obtained by Miceikaitė *et al*. demonstrating that FF estimator accuracy can decrease at low sequencing coverages (20). Severe FF estimation inaccuracies can lead to cases where a studied sample with a very low FF (which should fail the FF-based quality control) would be analysed, and the subsequent aneuploidy/euploidy estimation would be incorrectly considered as a confident estimate.

Although not comprehensively tested and analysed, we also observed that when several computational NIPT tools would be used as an ensemble (to supplement the sensitivity in different conditions), then for samples with FF ≥ 7%, such approach would provide perfectly accurate aneuploidy detection at 5M RPS (**Figure 4**) and also higher accuracy at 2.5M RPS as compared to the opposite strategy of using a single ‘best’ NIPT software tool.

While we had enough NIPT samples to compose a representative NIPT reference panel, we had a limited number of clinically validated trisomy samples to use. Although there is no reason to expect that adding trisomy samples to a given setup will alter the accuracy patterns of applied tools, future research could benefit from a comparison with a greater number of clinically validated trisomy samples, ideally sequenced at lower coverages. Although we tried to eliminate all possible sources of technical biases, as the artificial downsampling of samples (to lower sequencing read count per sample) might not perfectly reflect the natural read placement variability occurring in samples truly sequenced at lower coverages, these outcomes might possibly vary slightly. That said, a very similar downsampling approach has also been successfully implemented by Miceikaitė *et al*. to investigate fetal DNA fraction estimation accuracy at lower sequencing coverages (20). Our read placement variability effect analysis on Z-score inference was carried out only with the NIPTeR software, which nevertheless, was one of the most accurate tools in our comparison. In research and clinical practice utilising WGS-based NIPT with low sequencing coverage, similar analyses should be considered with any algorithm used for trisomy detection. A similar actual (i.e. non-downsampled) sequencing data based analyses or simulation can be carried out with any aneuploidy detection algorithm to assess the exact magnitude of this phenomenon.

This study provides further insights into WGS-based NIPT accuracy related to the experiment and the usage of different computational tools. Our results emphasise that confident trisomy detection is mainly affected by the computational tool selected for NIPT analysis in combination with sufficient sequencing coverage and accurately estimated fetal DNA fraction, as the currently available NIPT tools behave somewhat differently in the case of very low sequencing coverages.

## Supplementary data

**Supplementary Table 1.** Sample information for each of the analysed (subsample) group.

| **M RPS** | **Sample Group** | **SD** | **MIN** | **MEAN** | **MAX** | **N** |
| --- | --- | --- | --- | --- | --- | --- |
| 20 | T21 (clinically validated) | 4 277 096 | 14 347 516 | 24 163 449 | 32 626 785 | 19 |
| 15 | T21 (clinically validated) | 3 148 835 | 10 937 064 | 18 083 151 | 24 334 564 | 19 |
| 10 | T21 (clinically validated) | 2 081 671 | 7 209 906 | 11 958 958 | 16 127 678 | 19 |
| 5 | T21 (clinically validated) | 1 048 024 | 3 670 427 | 6 023 952 | 8 173 000 | 19 |
| 2.5 | T21 (clinically validated) | 523 966 | 1 833 823 | 3 011 415 | 4 086 277 | 19 |
| 1.25 | T21 (clinically validated) | 262 260 | 915 450 | 1 505 824 | 2 042 374 | 19 |
| 20 | T18 (clinically validated) | 2 791 633 | 18 450 798 | 23 124 624 | 27 336 184 | 8 |
| 15 | T18 (clinically validated) | 2 071 817 | 14 024 066 | 17 350 621 | 20 485 890 | 8 |
| 10 | T18 (clinically validated) | 1 395 351 | 9 235 616 | 11 476 668 | 13 625 081 | 8 |
| 5 | T18 (clinically validated) | 697 871 | 4 685 815 | 5 794 650 | 6 891 991 | 8 |
| 2.5 | T18 (clinically validated) | 348 455 | 2 344 985 | 2 898 169 | 3 446 589 | 8 |
| 1.25 | T18 (clinically validated) | 174 390 | 1 172 245 | 1 449 377 | 1 722 745 | 8 |
| 20 | T13 (clinically validated) | 2 980 546 | 19 042 280 | 22 183 555 | 24 971 957 | 3 |
| 15 | T13 (clinically validated) | 2 257 362 | 14 500 538 | 16 725 422 | 19 013 917 | 3 |
| 10 | T13 (clinically validated) | 1 469 534 | 9 567 867 | 11 039 287 | 12 506 927 | 3 |
| 5 | T13 (clinically validated) | 742 824 | 4 870 521 | 5 591 835 | 6 354 446 | 3 |
| 2.5 | T13 (clinically validated) | 370 783 | 2 435 984 | 2 796 437 | 3 176 752 | 3 |
| 1.25 | T13 (clinically validated) | 184 533 | 1 219 133 | 1 398 883 | 1 587 854 | 3 |
| 20 | Validation Low Risk | 3 594 972 | 13 643 477 | 23 747 773 | 32 396 033 | 393 |
| 15 | Validation Low Risk | 2 688 192 | 10 345 586 | 17 800 264 | 24 374 332 | 393 |
| 10 | Validation Low Risk | 1 791 357 | 6 839 734 | 11 799 169 | 16 181 461 | 393 |
| 5 | Validation Low Risk | 896 917 | 3 453 849 | 5 944 443 | 8 061 718 | 393 |
| 2.5 | Validation Low Risk | 448 456 | 1 726 568 | 2 972 178 | 4 028 665 | 393 |
| 1.25 | Validation Low Risk | 224 239 | 863 244 | 1 486 192 | 2 014 776 | 393 |
| 20 | Reference | 6 032 848 | 11 534 450 | 23 506 647 | 55 026 706 | 669 |
| 15 | Reference | 4 531 992 | 8 656 925 | 17 654 442 | 41 383 148 | 669 |
| 10 | Reference | 3 013 960 | 5 736 648 | 11 695 518 | 27 432 704 | 669 |
| 5 | Reference | 1 516 731 | 2 919 971 | 5 942 981 | 13 920 933 | 669 |
| 2.5 | Reference | 758 325 | 1 458 860 | 2 971 523 | 6 957 681 | 669 |
| 1.25 | Reference | 379 135 | 729 274 | 1 485 737 | 3 476 940 | 669 |

**Supplementary Table 2.** The number of false-negative and positive trisomy cases (with percentage) for each analysed software.

| **M RPS** | **Metric** | **WisecondorX** | **RAPIDR** | **NIPTmer** | **NIPTeR NCV** | **NIPTeR** | **GIPseq** |
| --- | --- | --- | --- | --- | --- | --- | --- |
| 20 | T21 FN | 0 (0) | 3 (0.735) | 0 (0) | 0 (0) | 0 (0) | 0 (0) |
| 20 | T21 FP | 1 (0.243) | 16 (3.922) | 1 (0.243) | 2 (0.485) | 1 (0.243) | 1 (0.245) |
| 15 | T21 FN | 0 (0) | 3 (0.732) | 0 (0) | 0 (0) | 0 (0) | 0 (0) |
| 15 | T21 FP | 0 (0) | 9 (2.195) | 0 (0) | 1 (0.243) | 1 (0.243) | 0 (0) |
| 10 | T21 FN | 0 (0) | 4 (0.973) | 2 (0.485) | 0 (0) | 0 (0) | 0 (0) |
| 10 | T21 FP | 1 (0.243) | 10 (2.433) | 1 (0.243) | 1 (0.243) | 2 (0.485) | 0 (0) |
| 5 | T21 FN | 0 (0) | 4 (0.971) | 2 (0.485) | 0 (0) | 0 (0) | 0 (0) |
| 5 | T21 FP | 0 (0) | 8 (1.942) | 0 (0) | 0 (0) | 0 (0) | 0 (0) |
| 2.5 | T21 FN | 2 (0.485) | 6 (1.456) | 5 (1.214) | 2 (0.485) | 2 (0.485) | 1 (0.243) |
| 2.5 | T21 FP | 0 (0) | 3 (0.728) | 0 (0) | 0 (0) | 0 (0) | 0 (0) |
| 1.25 | T21 FN | 5 (1.214) | 16 (3.883) | 9 (2.184) | 4 (0.971) | 4 (0.971) | 3 (0.728) |
| 1.25 | T21 FP | 1 (0.243) | 0 (0) | 1 (0.243) | 0 (0) | 0 (0) | 1 (0.243) |
| 20 | T18 FN | 0 (0) | 8 (2.015) | 0 (0) | 0 (0) | 0 (0) | 0 (0) |
| 20 | T18 FP | 1 (0.249) | 0 (0) | 0 (0) | 0 (0) | NA | 0 (0) |
| 15 | T18 FN | 0 (0) | 8 (2.005) | 1 (0.249) | 0 (0) | 0 (0) | 0 (0) |
| 15 | T18 FP | 2 (0.499) | 0 (0) | 0 (0) | 0 (0) | 0 (0) | 0 (0) |
| 10 | T18 FN | 0 (0) | 8 (2) | 1 (0.249) | 0 (0) | 0 (0) | 0 (0) |
| 10 | T18 FP | 1 (0.249) | 0 (0) | 0 (0) | 0 (0) | 0 (0) | 0 (0) |
| 5 | T18 FN | 0 (0) | 8 (1.995) | 1 (0.249) | 0 (0) | 1 (0.249) | 1 (0.249) |
| 5 | T18 FP | 0 (0) | 1 (0.249) | 0 (0) | 0 (0) | 0 (0) | 0 (0) |
| 2.5 | T18 FN | 1 (0.249) | 8 (1.995) | 3 (0.748) | 3 (0.748) | 3 (0.748) | 1 (0.249) |
| 2.5 | T18 FP | 0 (0) | 0 (0) | 0 (0) | 0 (0) | 0 (0) | 0 (0) |
| 1.25 | T18 FN | 3 (0.748) | 8 (1.995) | 5 (1.247) | 3 (0.748) | 4 (0.998) | 2 (0.499) |
| 1.25 | T18 FP | 0 (0) | 0 (0) | 1 (0.249) | 0 (0) | 0 (0) | 1 (0.249) |

**Supplementary Figure 1.**
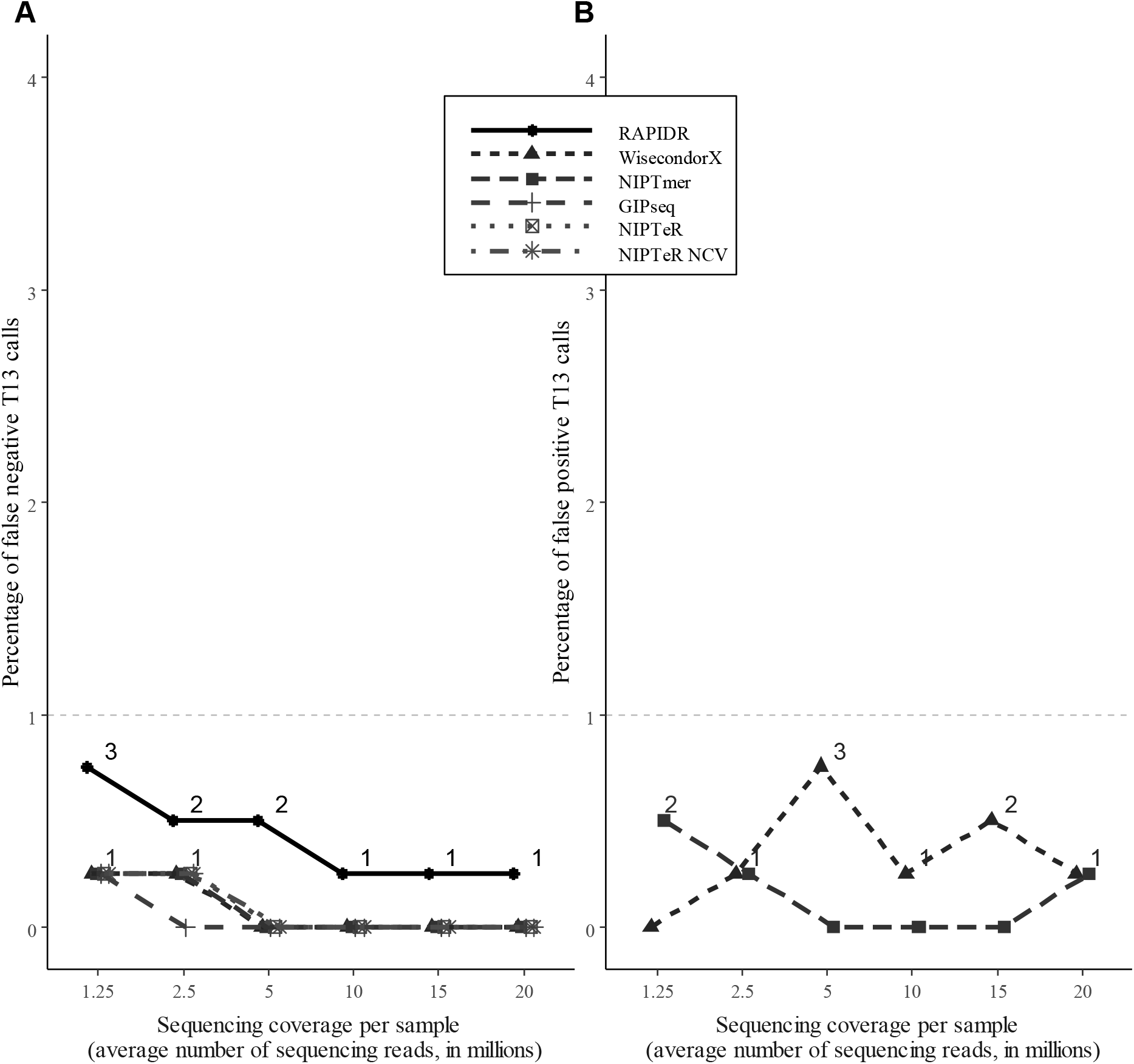
The percentage of false-negative (A) and false-positive (B) chromosome 13 trisomy cases in the case of different sequencing coverages.

**Supplementary Figure 2.**
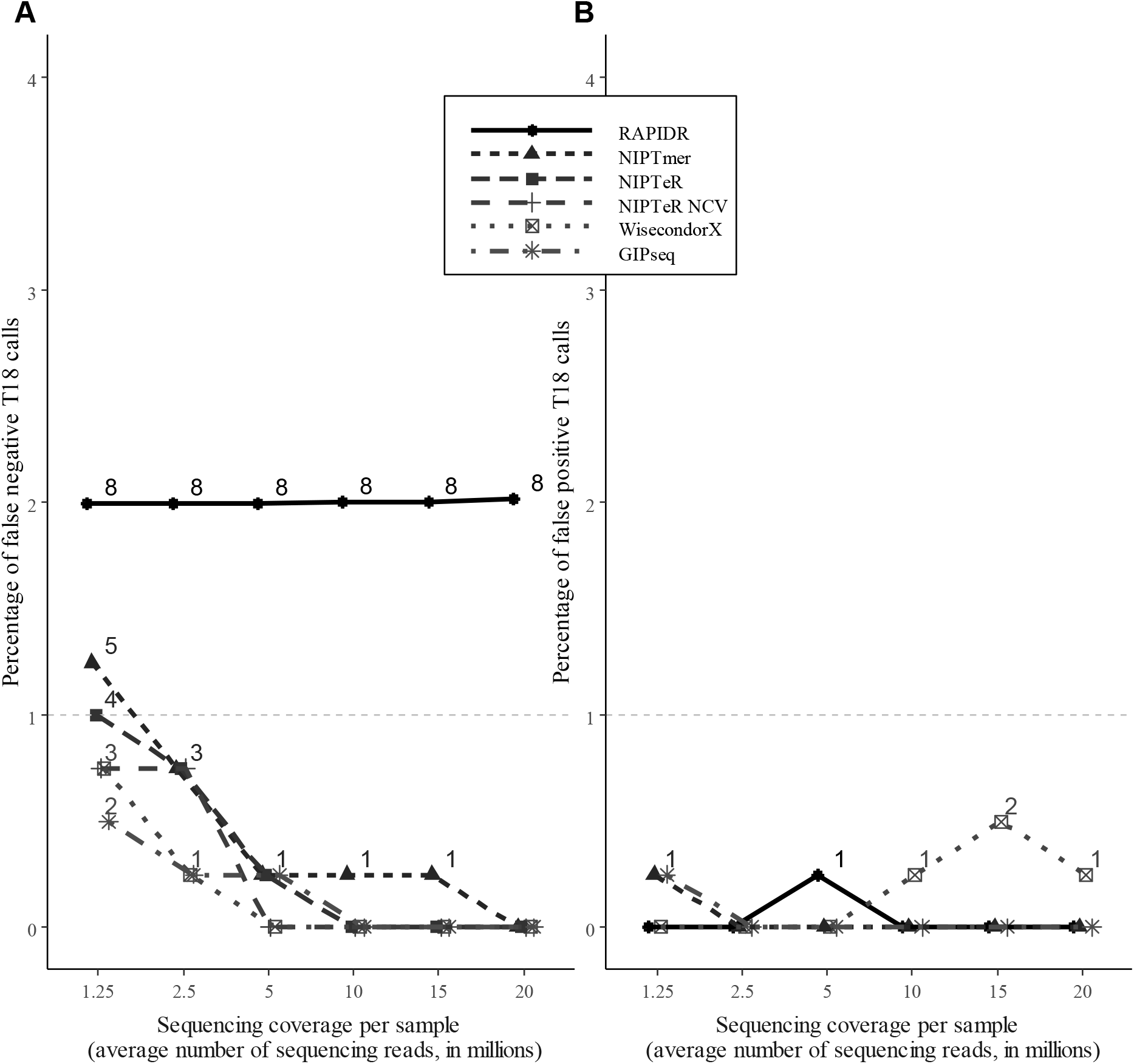
The percentage of false-negative (A) and false-positive (B) chromosome 13 trisomy cases in the case of different sequencing coverages.

**Supplementary Figure 3.**
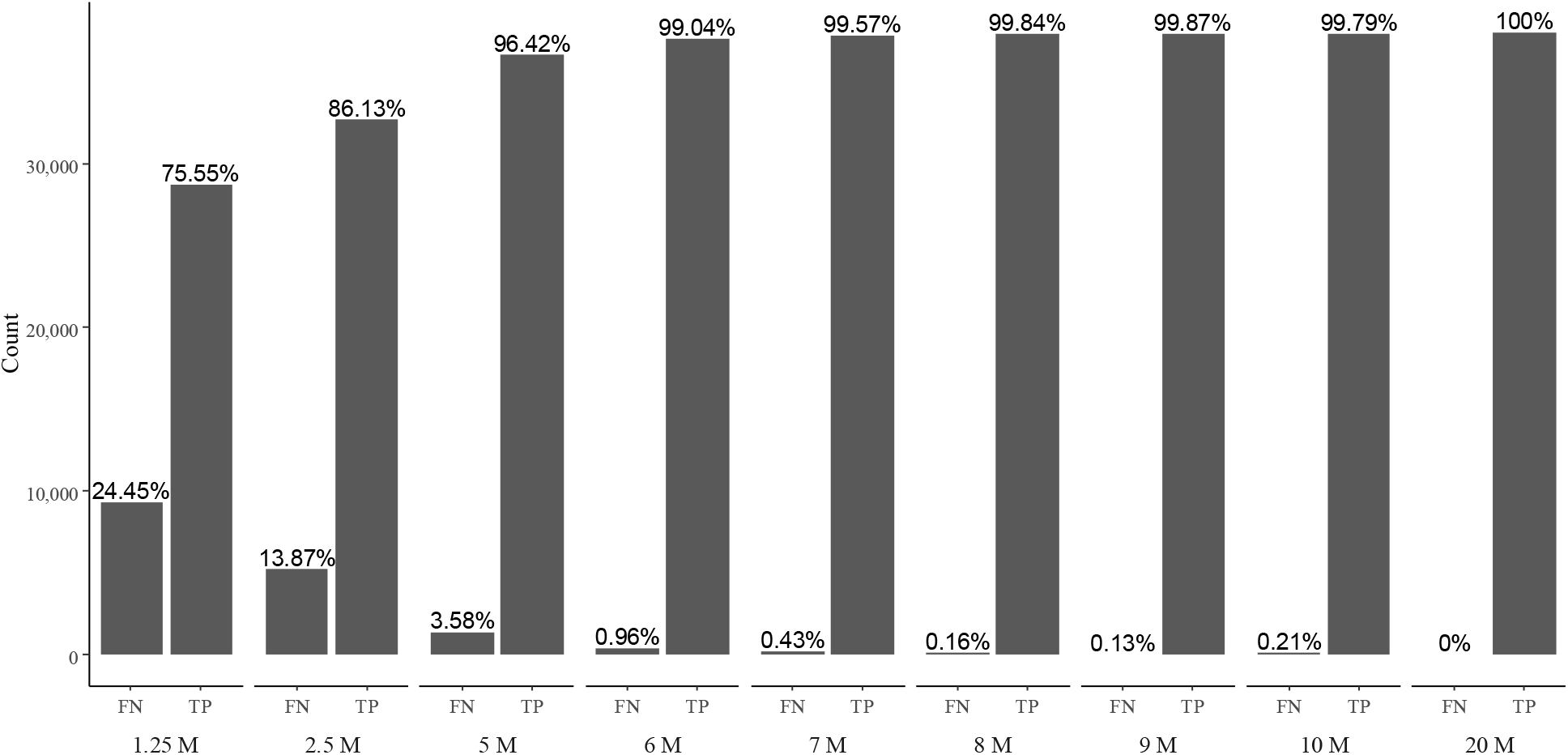
The effect of arbitrary (naturally occurring) sequencing read placement driven uncertainty on trisomy 21 inference. The natural sequencing read placement effect on T21 inference is significant with coverages lower than 7M RPS (0.96–24.5% of FN). With coverages 7M RPS and higher, the effect is insignificant (leads to less than 0.25% of FN T21 cases).

**Supplementary Figure 4.**
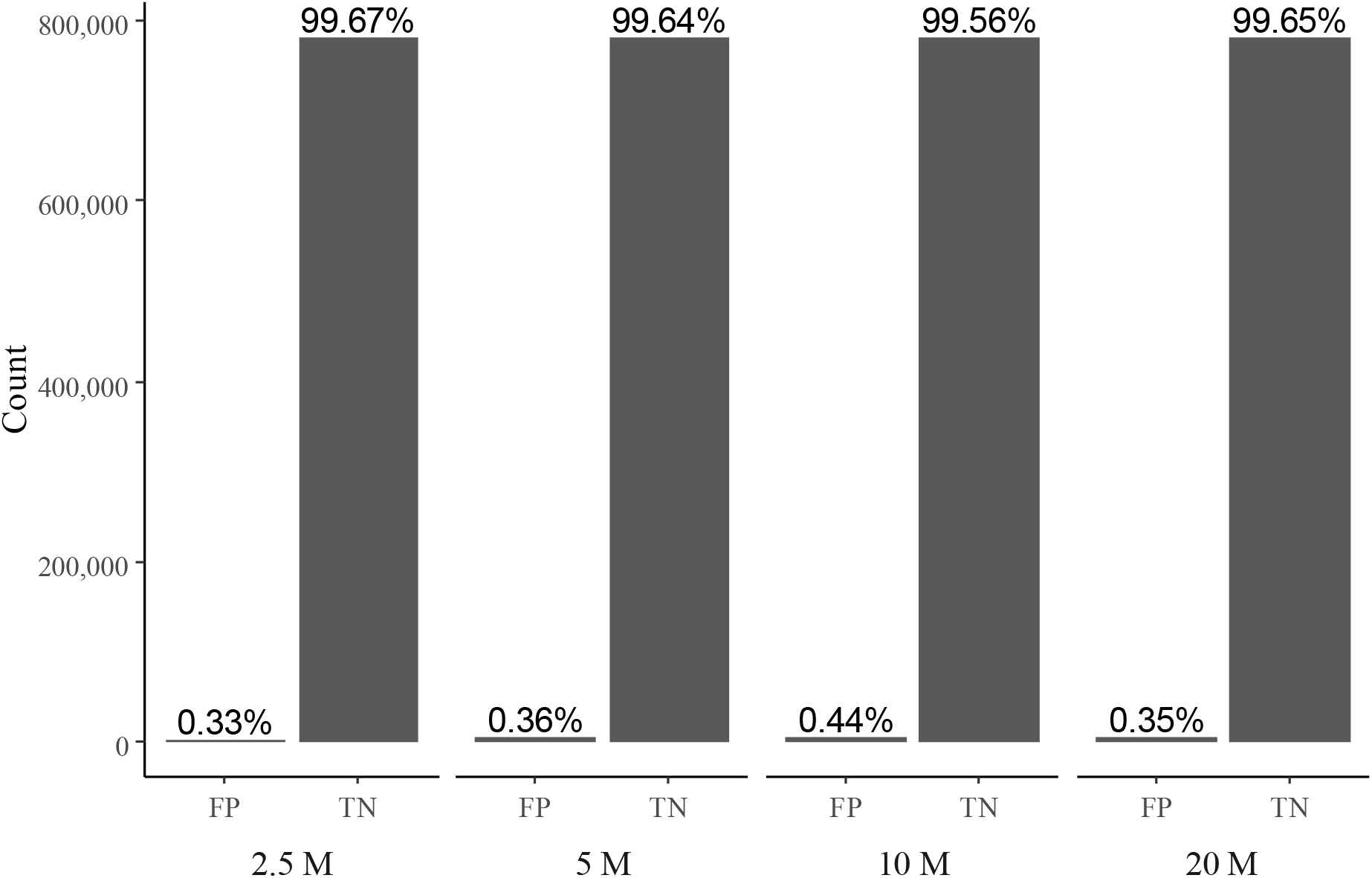
The effect of arbitrary (naturally occurring) sequencing read placement driven uncertainty on euploid sample calling and the resulting proportion of false-positive trisomy 21 cases. To summarise, 0.35%–0.44% of Z-scores depending on the subsample group were detected as FP T21.

**Supplementary Figure 5.**
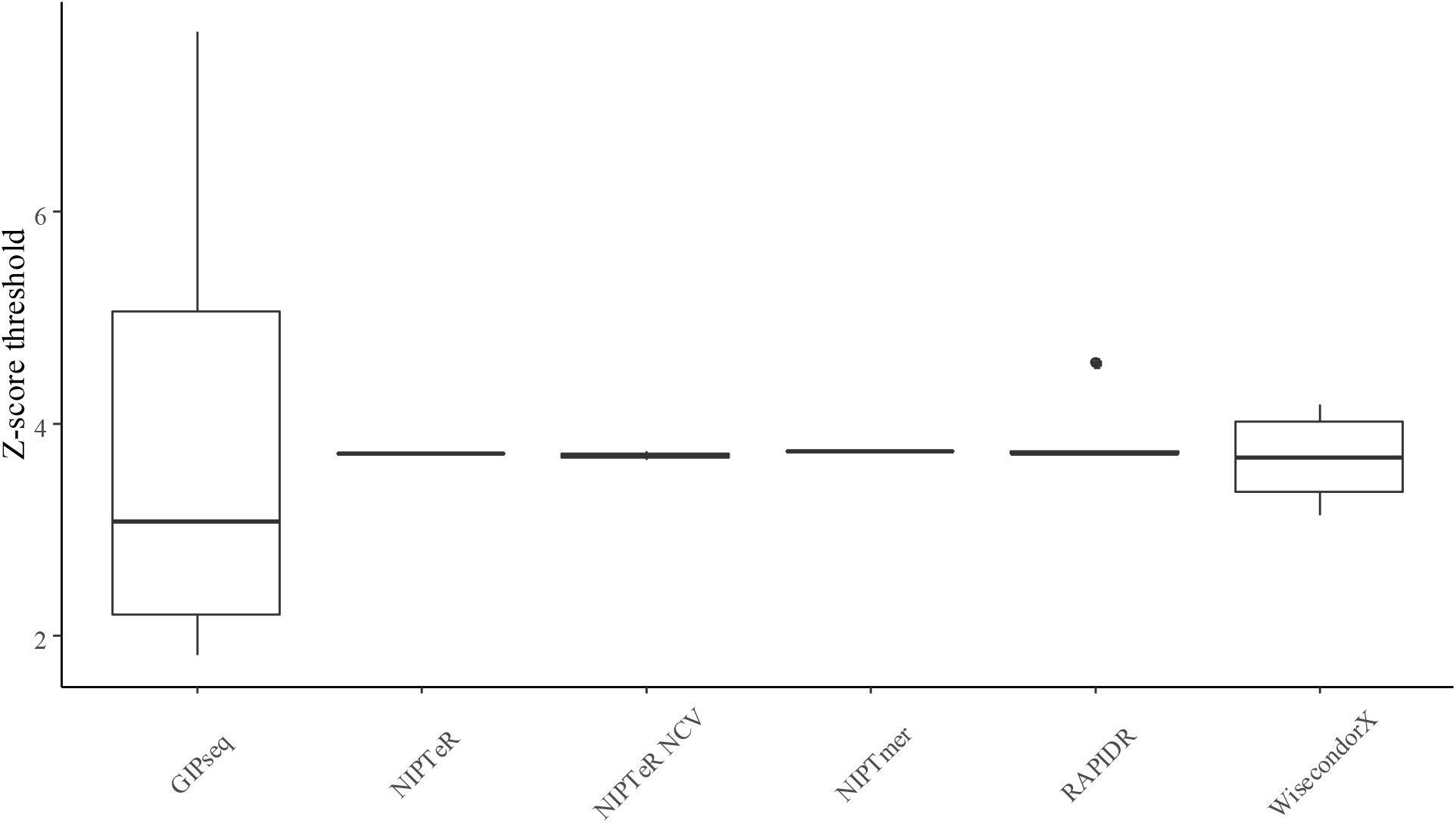
Variation in chromosome 21 *Z*_*e*_ cut-off thresholds on different sequencing depths in case of compared software tools.

**Supplementary Figure 6.**
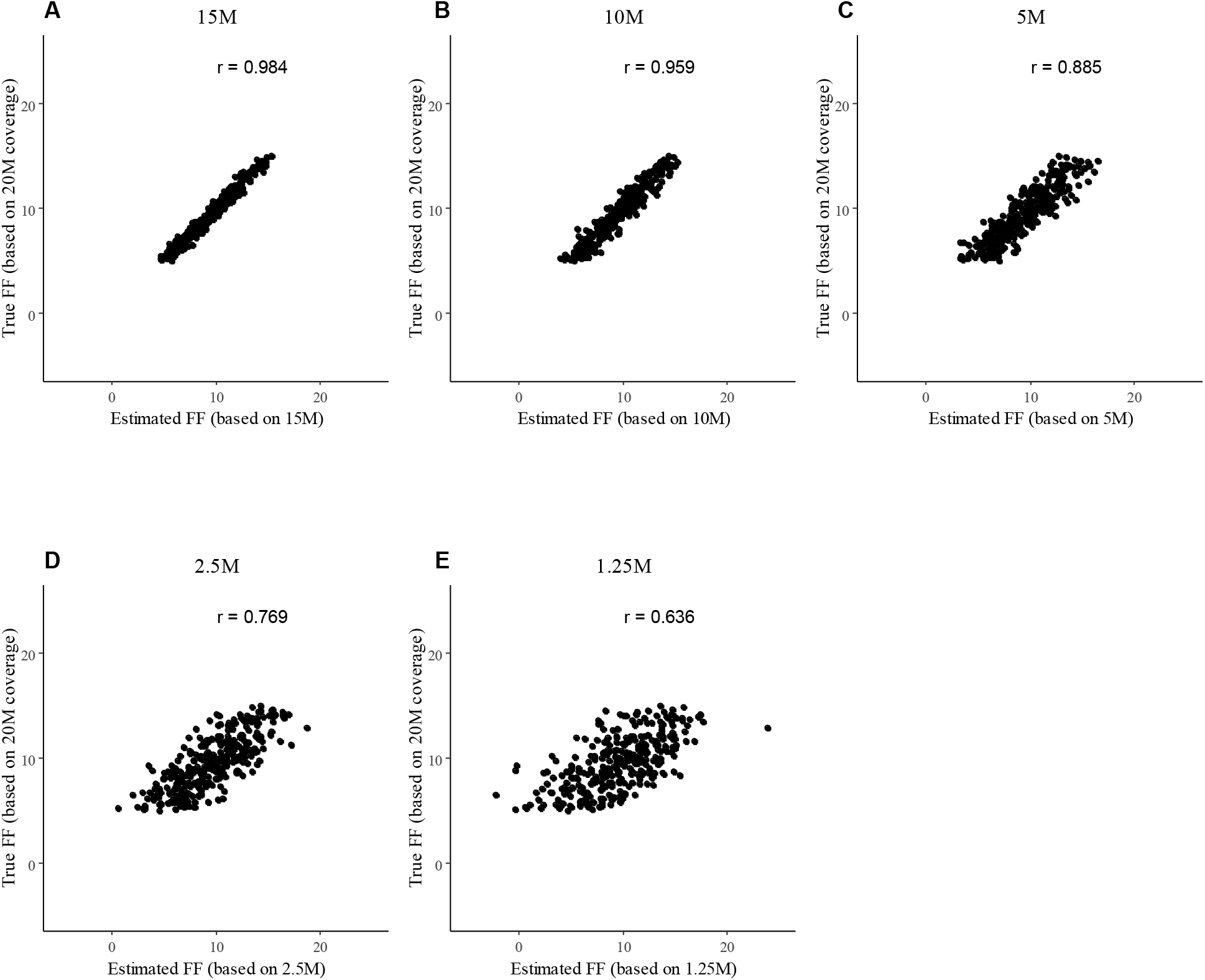
Comparison of 20M RPS FF estimates with the FF estimates at lower coverages. The Pearson correlation of the 20M RPS FF 5–15% estimates and the estimates on sequencing depths of 15M RPS (a), 10M RPS (b), 5M RPS (c), 2.5M RPS (d), and 1.25M RPS (e).

**Supplementary Figure 7.**
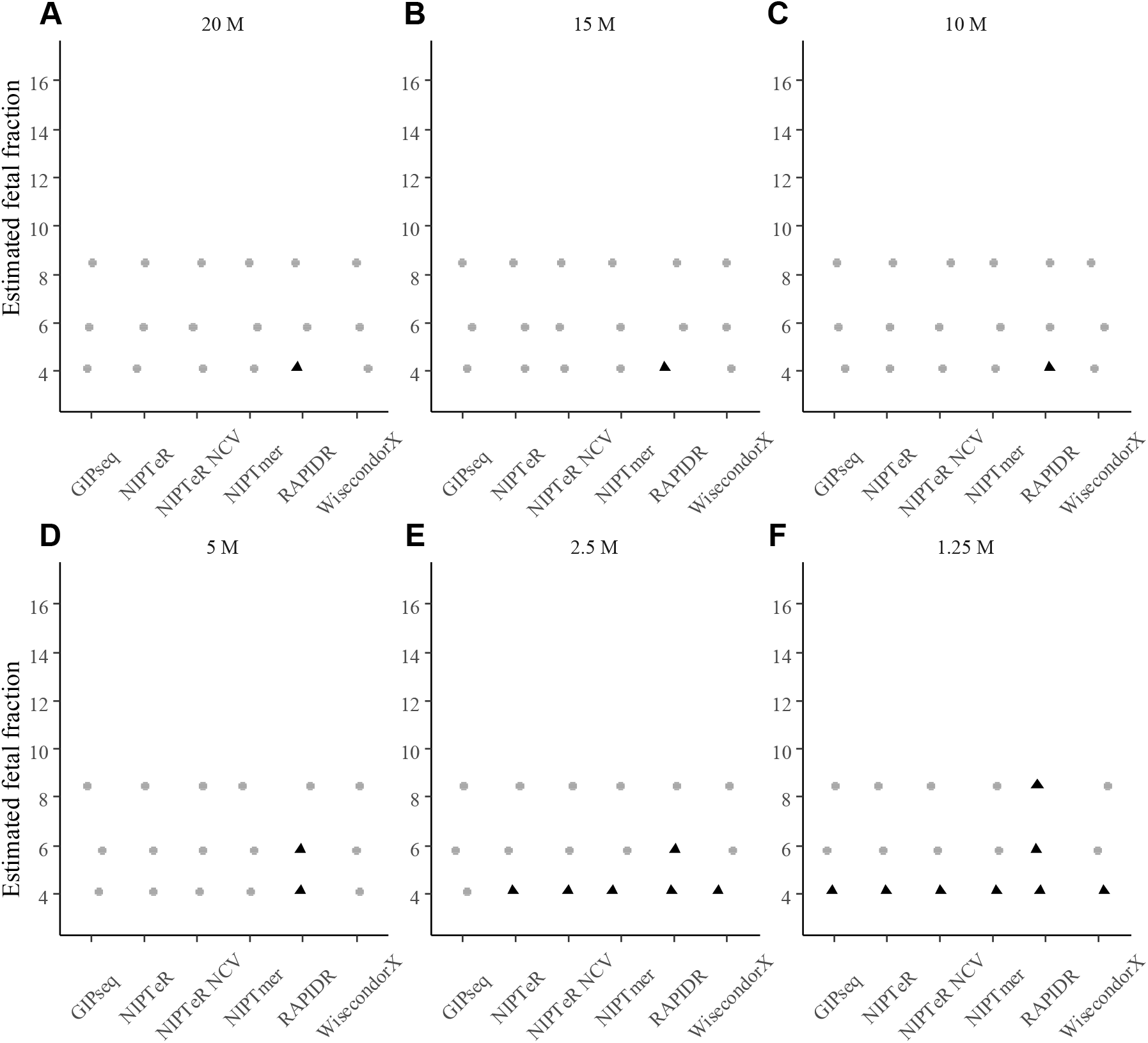
Fetal fraction effect on false-negative calls the of clinically validated trisomy 13 on different sequencing coverages. Computational tools were evaluated on 20 (A), 15 (B), 10 (C), 5 (D), 2.5 (E) and 1.25M RPS (F). The empirical cut-off was used for calling aneuploidy (internal classification in the case of GIPseq). Visualised samples are clinically validated T13 samples emulated to different coverages, and black triangles represent undetected aneuploidy. GIPseq was the only evaluated computational NIPT tool, which detected all three T13 samples at 2.5M RPS.

**Supplementary Figure 8.**
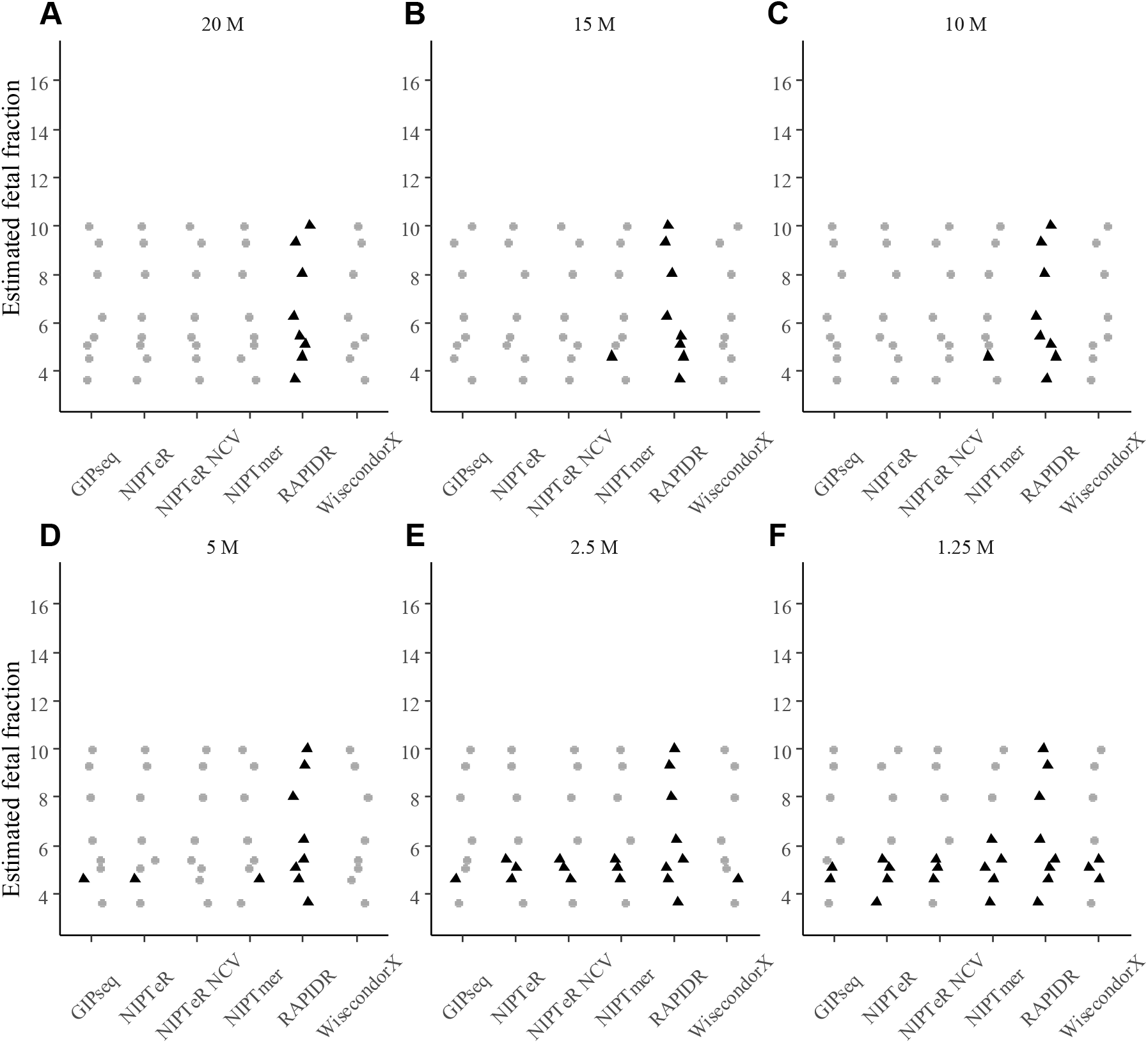
Fetal fraction effect on false-negative calls of the clinically validated trisomy 18 on different sequencing coverages. Computational tools were evaluated on 20 (A), 15 (B), 10 (C), 5 (D), 2.5 (E) and 1.25M RPS (F). The empirical cut-off was used for calling aneuploidy (internal classification in the case of GIPseq). Visualised samples are clinically validated T18 samples emulated to different coverages, and black triangles represent undetected trisomy.

## Acknowledgements

This work was supported by the Estonian Ministry of Education and Research [IUT34-16]; and the Enterprise Estonia [EU53935, EU48695]. The presented NIPT computational analyses were carried out in the High-Performance Computing Center of the University of Tartu.

